# Impairment of mitochondrial respiratory function as an early biomarker of apoptosis induced by growth factor removal

**DOI:** 10.1101/151480

**Authors:** Hélène Lemieux, Patrick Subarsky, Christine Doblander, Martin Wurm, Jakob Troppmair, Erich Gnaiger

## Abstract

Intracellular signaling pathways not only control cell proliferation and survival, but also regulate the provision of cellular energy and building blocks through mitochondrial and non-mitochondrial metabolism. Wild-type and oncogenic RAF kinases have been shown to prevent apoptosis following the removal of interleukin 3 (IL-3) from mouse pro-myeloid 32D cells by reducing mitochondrial reactive oxygen species production. To study primary effects of RAF on mitochondrial energy metabolism, we applied high-resolution respirometry after short-term IL-3 deprivation (8 h), before 32D cells show detectable signs of cell death. Respiration in intact 32D cells was suppressed as an early event following removal of IL-3, but remained more stable in 32D cells expressing the v-RAF oncogene. In permeabilized 32D cells deprived of IL-3, respiratory capacities of the NADH-pathway, the convergent NADH&succinate-pathway, and Complex IV activity were decreased compared to cells grown in the presence of IL-3, whereas succinate-supported respiration remained unchanged, consistent with control by Complex IV. The apparent Complex IV excess capacity was zero above NADH&succinate-pathway capacity reconstituting tricarboxylic acid cycle function. In comparison, electron flow reached only 60% when supported by succinate alone through Complexes II, III and IV, and was therefore relatively insensitive to Complex IV injuries up to a threshold of 40% inhibition. A slight increase in respiration following addition of cytochrome *c*, a marker of mitochondrial outer membrane leakage, was present after IL-3 depletion, indicating mitochondrial fragility. Our results highlight a novel link between the key mitogenic and survival kinase CRAF and mitochondrial energy homeostasis.

## Introduction

RAF serine-threonine kinases are evolutionary highly conserved components of cellular signal transduction, which promote cell growth, transformation and survival ^1, 2^. The RAF kinase family comprises the originally identified viral oncogene v-RAF and the subsequently cloned proto-oncogenes CRAF, ARAF and BRAF ^3^. In melanoma, the activation of the mitogen-activated protein kinase pathway by BRAF mutants has been shown to be the primary cause of malignant transformation ^4^.

The anti-apoptotic function of CRAF is well documented by overexpression and gene knockout studies ^5^. Cell death is increased in CRAF and BRAF deficient animals ^6-11^. Treatment with BRAF inhibitors significantly improves the survival of patients with metastatic melanoma ^4^. The mitochondrion is a key organelle involved in the regulation of apoptosis. Studies on the control of cell survival by CRAF suggest a role for this kinase in maintaining mitochondrial (mt) integrity and counteracting induction of apoptosis (reviewed by ^5^). In vascular endothelial cells, fibroblast growth factor induces CRAF translocation and its binding to mitochondria through the N-terminal domain (reviewed by ^12^). Cells are protected from apoptosis when RAF accumulates in the mitochondria ^8, 13, 14^.

Crosstalk between mitochondria and other compartments of the cell has important regulatory functions and is mediated by small molecules, such as reactive oxygen species (ROS) and Ca^2+^, and by components of intracellular signaling pathways ^15-17^. Oncogenically activated signaling proteins are implicated in the metabolic rewiring of tumor cells including the well-established shift from oxidative phosphorylation to glycolysis under normoxic condition ^4, 18, 19^. Thus the frequently mutated RAS oncogene, an essential upstream component in the activation of RAF kinases, exerts many effects on mitochondria including an increase in mt-mass, mtDNA content ^20^, concentrations of tricarboxylic acid (TCA) cycle intermediates ^21^, and modifications of mt shape and distribution ^13^. Furthermore, RAS, located upstream of RAF protein in the RAS/RAF/MEK pathway, is linked to an increase in mt-production of ROS ^20, 22, 23^. K-RAS protein is localized to the mitochondria and causes mt-respiratory dysfunction at an early time point ^24^. In fact, more than a third of human tumors present an activated mutant RAS oncogene ^25, 26^.

Small GTPases of the RAS family have effectors other than RAFs. Some studies demonstrate increased ROS levels in cells transformed by RAS ^23^, whereas RAF-transformed cells upregulate defense mechanisms by increasing antioxidant capacity ^27, 28^. In hepatocellular carcinoma cells resistant to treatment by the multikinase inhibitor sorafenib also targeting RAF, activating oxidative phosphorylation (OXPHOS) by the pyruvate dehydrogenase kinase inhibitor dichloroacetate markedly sensitizes the cancer cells to apoptosis ^29, 30^. A similar observation has been reported for melanoma cells carrying the activating V600E mutation in BRAF ^18^. Mutant BRAF in melanoma cells downregulates expression of PGC1α, a member of the superfamily of transcriptional co-activators that promote mitochondrial biogenesis and enhance OXPHOS (see review ^4^). A recent study in pancreatic β cells provides further evidence that the increase in mitochondrial respiration triggered by PKC is in part mediated through CRAF ^31^. While these studies illustrate the profound role of RAF kinases in reprogramming tumor cell metabolism and preventing cell death, they do not address a possible direct effect of RAF on OXPHOS and bioenergetic coupling at earlier time points, which precede the onset of apoptotic cell death.

The goal of our study was to analyze mitochondrial respiratory function after a short absence of growth factor with the known consequences on the shutdown of intracellular signaling cascades, as it may occur *in vivo* during stroke or ischemia/reperfusion in solid organ transplantation. Focusing on an early time point, before any molecular signals of apoptosis can be detected, we demonstrated that the significant decline of mitochondrial respiratory capacity is the earliest elucidated effect of growth factor removal. Loss of mt-function was effectively suppressed by oncogenic RAF (v-RAF), suggesting for the first time a direct link between the key mitogenic and survival kinase CRAF and mitochondrial energy homeostasis.

## Results

### Cell survival and protein content are preserved at an early time point of growth factor withdrawal

IL-3 withdrawal from 32D cells (32D-IL3) provides a unique system for the study of cellular alterations that precede the onset of apoptotic cell death. Absence of growth factor for 24 h reproducibly results in apoptosis of 80% of the parental 32D cells and in only 20% of the 32D cells expressing v-RAF (32Dv-RAF cells) ^32^. Activated RAF prevents late mitochondrial events preceding cell death, e.g., increase in ROS and mitochondrial Ca^2+ 27, 28^. The present study focuses on the earliest changes following growth factor removal. IL-3 deprivation for 8 h resulted in no detectable signs of cell death (Fig. 1A). Re-addition of IL-3 at the end of the 8- h starvation period prevented subsequent cell death. In all our experiments, the starting cell viability was higher than 96% and remained unchanged after 8 h of IL-3 withdrawal in 32D and 32Dv-RAF cells. A significant decrease in cell volume was observed in 32D cells [0.4 (0.83- 1.37) pL versus 0.79 (0.69-0.93) pL] but not in v-RAF protected cells (Table 1). Regardless of the decrease in cell volume, the protein content of 32D and 32Dv-RAF cells was not significantly different after IL-3 withdrawal (Table 1). Furthermore, IL-3 withdrawal for 8 h did not cause detectable effects on cell cycle distribution (Table 2).

**Fig. 1:**
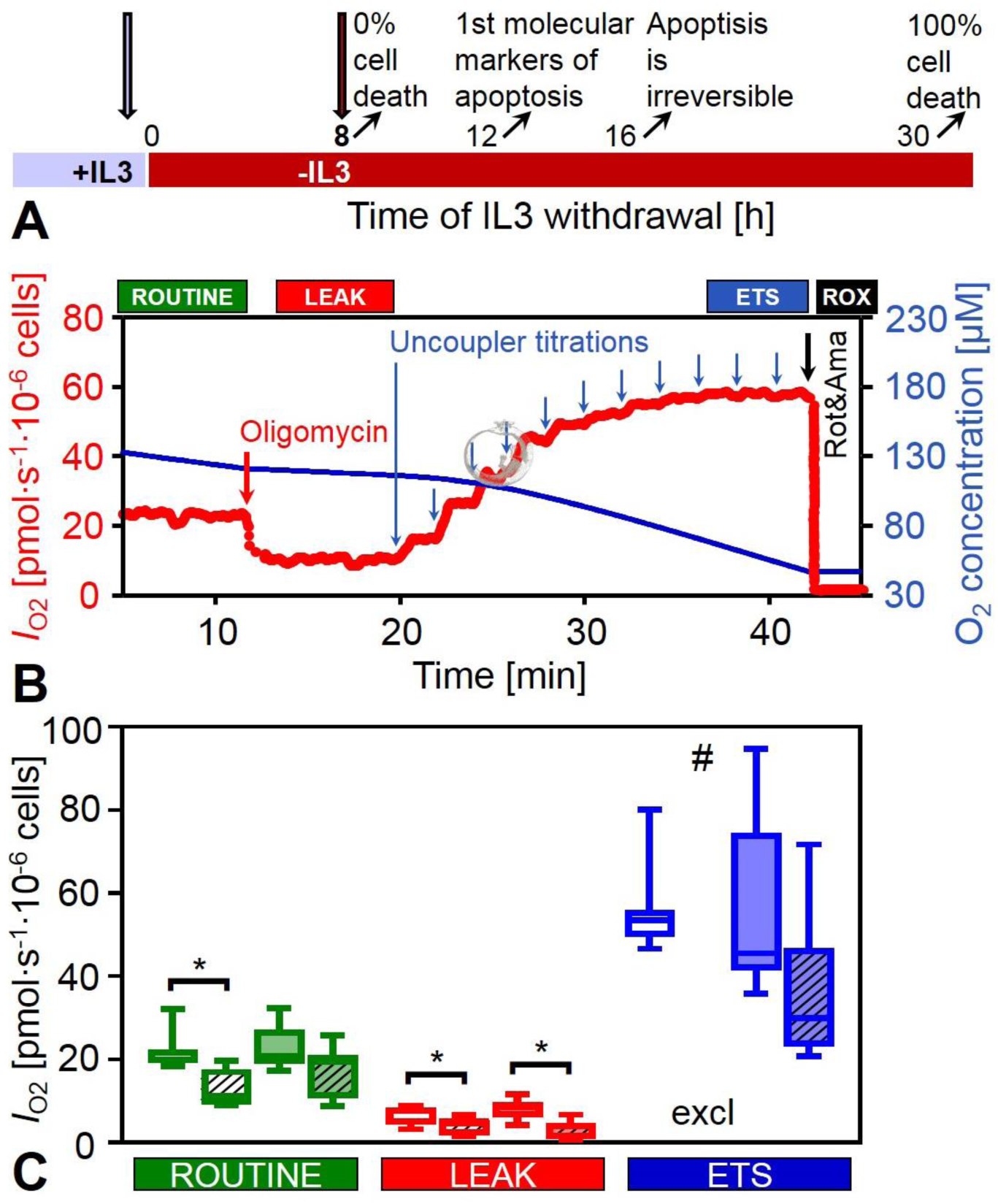
Effect of 8 h IL-3 withdrawal on respiration, *I*_O2_, of intact 32D and 32Dv-Raf cells. (A) Experimental design in the time course of IL3 withdrawal. (B) Representative trace of high-resolution respirometry in intact 32D cells in the presence of the growth factor IL-3. Oxygen consumption (left axis, bold line) and oxygen concentration (right axis, thin line) are plotted as a function of time. Arrows indicate times of titrations in a simple coupling control protocol. ROUTINE respiration is the activity of intact cells in culture medium (RPMI) containing exogenous substrates which support energy metabolism and biosynthesis. LEAK respiration is the minimum respiration compensating mainly for proton leak after inhibition of ATP synthase by olygomycin. ETS is the maximal electron transfer capacity at optimum uncoupler concentration. Complexes I and III were inhibited by rotenone and antimycin A for measurement of residual oxygen consumption (ROX). (C) Box plots indicate the minimum, 25^th^ percentile, median, 75^th^ percentile, and maximum (*n*=8 replicates; *N*=3 independent cultures). * Significantly different (*p*<0.05) after IL-3 withdrawal. # A latent injury was revealed in samples when uncoupler titration following oligomycin resulted in respiration not stimulated above the ROUTINE level. Since the actual ETS capacity cannot be lower than ROUTINE respiration, these data are excluded from the graph (excl).

**Table 1:** Cell size, protein content, citrate synthase (CS) and lactate dehydrogenase (LDH) activities in 32D and 32D v-Raf cells, grown in the presence of IL-3 or after 8 hours of IL-3 withdrawal. Median (min-max), *n*=4-5.

|  | 32D |  | 32D v-Raf |  |
| --- | --- | --- | --- | --- |
|  | +IL3 | -IL3 | +IL3 | -IL3 |
| Cell volume [pL] | 0.94 (0.83-1.37) | 0.79 (0.69-0.93)* | 1.06 (0.99-1.09) | 0.90 (0.84-1.16) |
| Protein content [mg·10<sup>-6</sup> cells] | 0.13 (0.06-0.14) | 0.17 (0.11-0.27) | 0.16 (0.15-0.21) | 0.15 (0.11-0.19) |
| CS activity [U·10<sup>-6</sup> cells] | 0.024 (0.021-0.030) | 0.021 (0.015-0.030) | 0.029 (0.022-0.045) | 0.025 (0.021-0.043) |
| LDH activity [U·10<sup>-6</sup> cells] | 0.029 (0.018-0.041) | 0.023 (0.018-0.038) | 0.037 (0.025-0.057) | 0.029 (0.023-0.055) |

\*Significantly different from the same cells in the presence of IL-3 (+IL3); $p<0.05$ .

**Table 2:** Effect of IL-3 starvation on cell cycle distribution of 32D and 32Dv-Raf cells grown in the presence of IL-3 or after 8 h IL-3 withdrawal. Values are means±SD (*n*=5) in percent of cells in each phase of the cycle.

| Cell cycle phase | 32D |  | 32D v-Raf |  |
| --- | --- | --- | --- | --- |
|  | +IL3 | -IL3 | +IL3 | -IL3 |
| Sub G0 | 14±4 | 14±7 | 6±3 | 6±3 |
| G1 | 53±3 | 55±4 | 54±1 | 55±2 |
| S | 8±3 | 8±5 | 13±4 | 13±2 |
| G2 | 21±2 | 19±4 | 23±2 | 22±3 |

### Activated RAF protects mitochondrial function during growth factor deprivation, while mitochondrial and glycolytic marker enzymes are preserved

ROUTINE respiration (*R*) of intact 32D cells was suppressed significantly after 8 h growth factor deprivation (Fig. 1). The decline was not significant in 32Dv-RAF cells (Fig. 1C), suggesting that the presence of activated RAF was sufficient to prevent this early change caused by IL-3 withdrawal. Resting oxygen consumption that compensates for proton leak, electron slip and cation cycling (LEAK respiration, *L*), was measured by inhibition of ATP-synthase with oligomycin (Fig. 1B). LEAK respiration was significantly suppressed in 32D cells after 8 h growth factor deprivation (Fig. 1C). The decrease in LEAK respiration following IL-3 removal, however, was not prevented by v-RAF overexpression. Electron transfer system (ETS) capacity (*E*) was estimated by uncoupling the cells with optimal concentrations of the protonophore FCCP (Fig. 1B). The apparent ETS excess capacity above ROUTINE respiration, *E-R*, was identical in 32D and 32-v-RAF cells (Fig. 1C). ETS capacity declined not significantly in 32-v-RAF cells after IL-3 removal. 32D-IL3 cells, however, presented a latent injury revealed by uncoupler titration after oligomycin treatment, when respiration was unstable and flux was occasionally not even stimulated to the level of ROUTINE respiration (data not shown). An appropriate estimation of ETS capacity was not possible in those cells after inhibition by oligomycin.

Activities of the mitochondrial marker enzyme citrate synthase (CS) and the glycolytic marker enzyme lactate dehydrogenase (LDH) expressed in units per million cells did not differ significantly between parental and v-RAF expressing 32D cells and were not significantly reduced by IL-3 deprivation (Table 1). Variations in the activities of these marker enzymes were directly correlated, suggesting no re-programming of glycolytic versus aerobic metabolism under the prevailing growth conditions (Fig. 2).

**Fig. 2:**
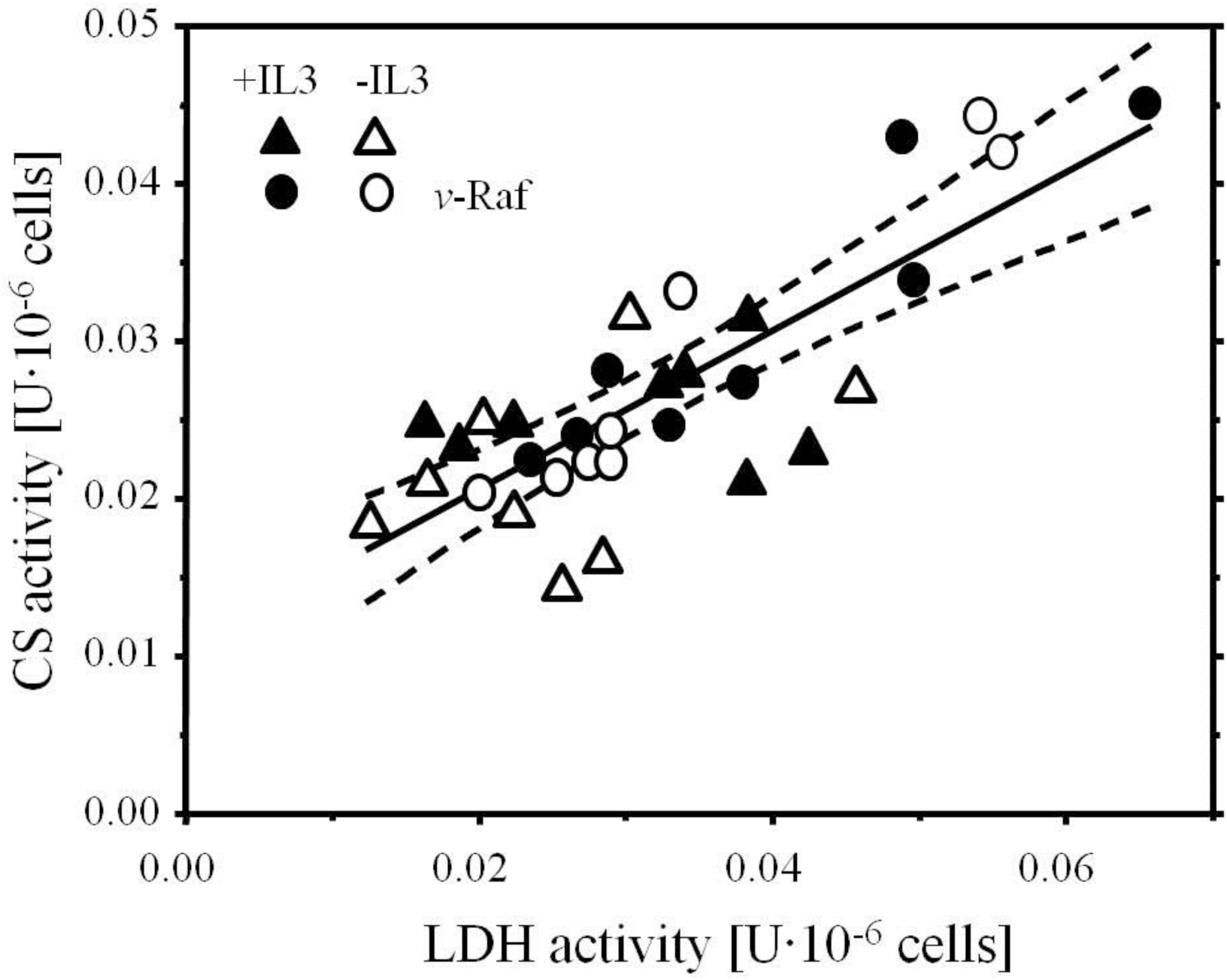
Correlation of the activities of citrate synthase (CS) and lactate dehydrogenase (LDH) expressed per million cells.

### Growth factor deprivation causes a defect in OXPHOS and ETS capacity and cytochrome c release

In order to assess specific defects that cause the decrease in mitochondrial respiratory function after IL-3 removal in the 32D cells, we applied OXPHOS analysis in cells permeabilized by digitonin. Cells treated with an apoptosis inducer show a change in the sensitivity of the outer mt-membrane to digitonin ^33^. Therefore, we tested permeabilization of the cell membrane by digitonin in 32D and 32Dv-RAF cells grown in the presence or absence of IL-3. Cells were incubated in mitochondrial respiration medium (MiR05) in the ROUTINE respiratory state supported by endogenous substrates. Rotenone passes the cell membrane, inhibits Complex I (CI), and inhibits respiration immediately. External succinate (S) and ADP are impermeable in intact cells and do not stimulate respiration. Digitonin permeabilizes the plasma membrane when titrated above a threshold concentration. Then succinate and ADP become accessible to the mitochondria and respiration is stimulated to the state of S- (CII-linked) OXPHOS capacity (Fig. 3). The digitonin concentration required for membrane permeabilization was identical in 32D and 32Dv-RAF cells. However, cell membrane permeabilization by digitonin was delayed in 32D-IL3 compared to 32D+IL3 cells, as shown by the slow increase in respiration of the 32D-IL3 cells (Fig. 3A, dashed line). In contrast, the time course of cell membrane permeabilization in 32Dv-RAF cells was identical after 8 h with or without growth factor (Fig. 3B).

**Fig. 3:**
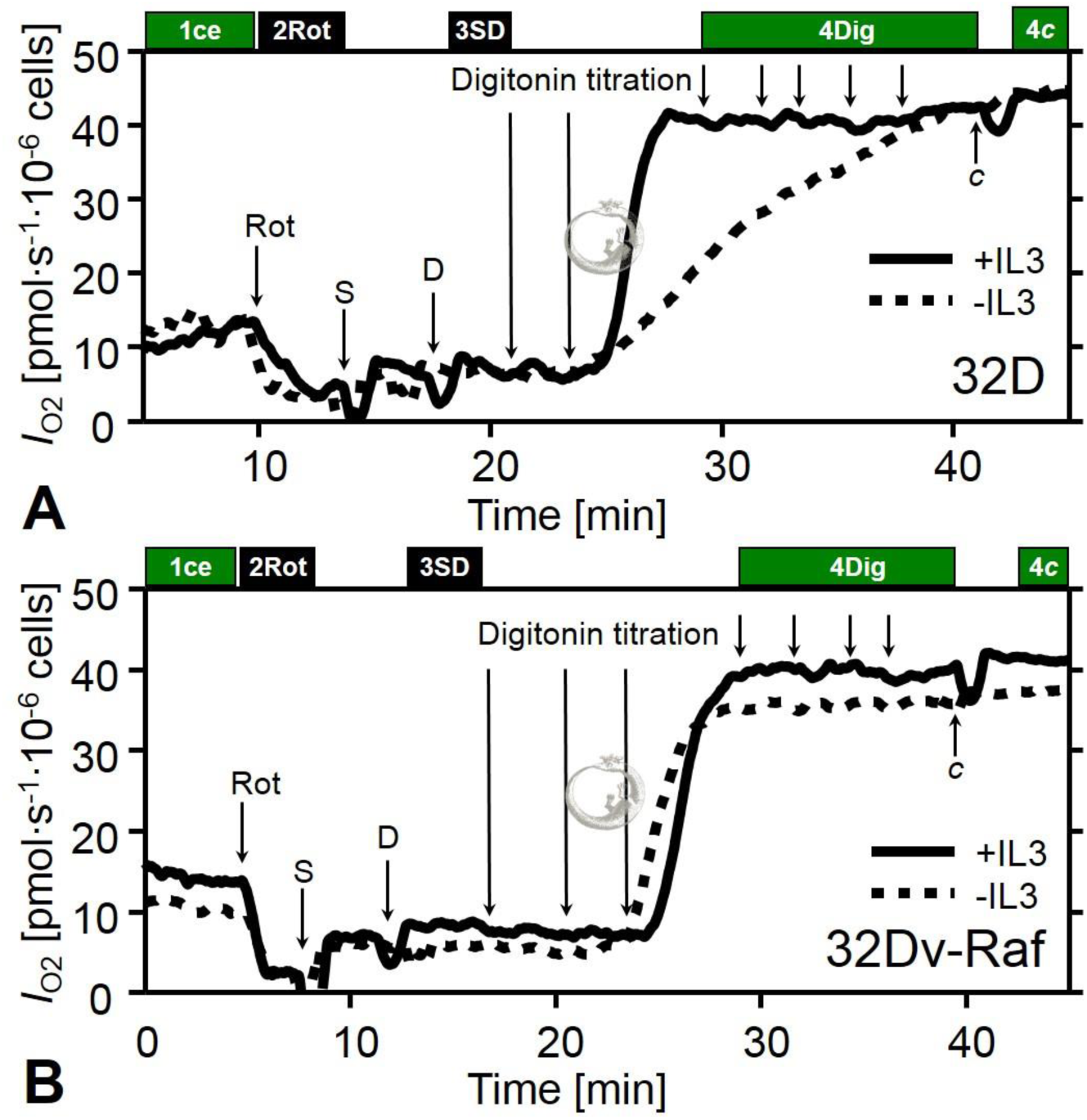
Optimization of digitonin concentration for selective permeabilization of the plasma membrane of 32D cells (A) and 32-v-RAF cells (B) in the presence of IL-3 (full lines) and after 8 h IL-3 withdrawal (dotted lines). Arrows indicate times of titrations. ROUTINE respiration was recorded after addition of intact cells (1 ce) in MiR05 medium. Rotenone (2Rot) inhibits CI, blocks formation of succinate in the TCA cycle and stops simultaneously electron transfer through CII. Thus respiration is inhibited almost completely to the level of residual oxygen consumption (ROX). S-pathway OXPHOS capacity is obtained after addition of succinate and ADP (3SD) and titrations of digitonin (4Dig; final concentration in the chamber: 1, 2, 3, 4, 6, 10 and 20 μg·ml^−1^). 10 μM cytochrome *c* (4*c*) did not significantly stimulate respiration, indicating that the outer mt-membrane remained intact even at the highest digitonin concentration.

Following optimization of the permeabilization conditions we performed high-resolution respirometry with the objective to diagnose the specific defects of the OXPHOS system caused by IL-3 withdrawal in 32D cells. The substrate-uncoupler-inhibitor titration (SUIT) protocol is shown in Fig. 4A and B. After measurement of ROUTINE respiration of intact cells in MIR05, the plasma membrane was permeabilized by digitonin, and NADH-linked substrates (supporting the N-pathway; pyruvate, glutamate and malate, PGM) were added to measure LEAK respiration in the absence of adenylates. OXPHOS capacity (*P*) was measured after addition of a saturating concentration of ADP (coupling control). NADH-linked LEAK respiration and OXPHOS capacity were reduced significantly in 32D-IL3 cells compared to controls (Fig. 4C). Cytochrome *c* release was evaluated indirectly by addition of cytochrome *c* (Fig. 4B). No significant stimulation by cytochrome *c* was observed in 32D+IL3 cells. In contrast, the decrease of respiration in 32D-IL3 cells was associated with a small but significant increase in respiration after cytochrome *c* addition. The cytochrome *c* control factor, *FCFc* = 1- PGM/PGM*c*, was 0.15 (0.04-0.24), significantly different from zero (paired t-test). After further addition of succinate to support convergent NADH&succinate-linked (NS) electron flow through Complexes I&II into the Q-cycle (PGMS-pathway control), respiration was increased approximately two-fold. The difference between 32D-IL3 and 32D+IL3 cells remained significant (Fig. 4C).

**Fig. 4:**
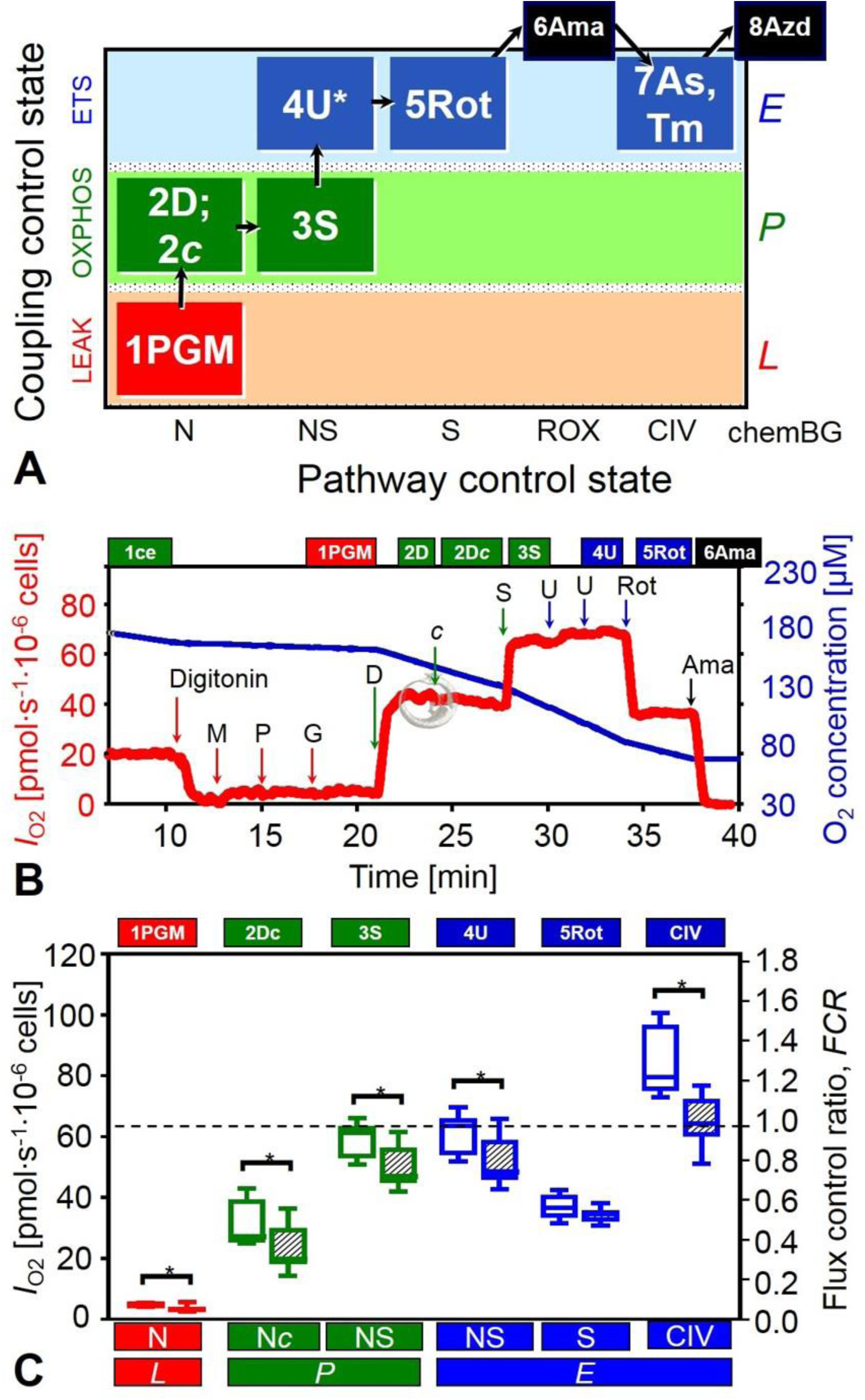
Effect of IL-3 withdrawal on mitochondrial respiration in digitonin permeabilized 32D cells. In the substrate-uncoupler-inhibitor-titration (SUIT) protocol, LEAK respiration was estimated in the presence of the NADH-linked (N) substrates pyruvate (P), malate (M) and glutamate (G) in the absence of adenylates. OXPHOS capacity (N-pathway; PGM*_P_*) was measured after addition of ADP. Cytochrome *c* (*c*) was added to test the integrity of the outer mitochondrial membrane (PGM*c_P_*). Subsequent succinate (S) addition allowed the measurement of respiration with convergent NS-electron flow through Complexes I&II (PGMS*_P_*). Electron system capacity (ETS) was evaluated by uncoupling with optimum FCCP concentration (PGMS*_E_*). Succinate-supported respiration through Complex II (S) was obtained by inhibition of CI with rotenone (Rot). Inhibition of Complex III by antimycin A (Ama) provided an estimation of residual oxygen consumption (ROX). Cytochrome *c* oxydase activity (Complex IV, CIV) was measured by addition of ascorbate and TMPD and correction of chemical background of autooxidation. (A) Coupling/pathway control diagram of the SUIT protocol. The asterisk (4U*) indicates multiple uncoupler titrations. (B) Representative experiment showing oxygen flow, *I*_O2_ (thick line), and oxygen concentration (thin line). Respiration is expressed in pmol O2 per second per million cells. Arrows indicate the times of titrations. Cellular ROUTINE respiration was measured in respiration medium (MIR05) in the absence of exogenous substrates after addition of intact cells (1ce). Cells were permeabilized with optimum digitonin concentration (10 μg/ml). (C) Box plots indicate the minimum, 25 ^th^ percentile, median, 75^th^ percentile, and maximum (*n*=12 replicates; *N*=3 independent cultures). * significantly different (*p*<0.05) after IL-3 withdrawal.

### Succinate pathway capacity and coupling are preserved during growth factor deprivation

Uncoupler titrations were performed in the ADP-activated state with PGMS, to evaluate ETS capacity (*E;* noncoupled) in relation to OXPHOS capacity (*P*; coupled; Fig. 4C). An excess *E-P* capacity factor, 1-*P/E*, above 0.0 indicates limitation of OXPHOS capacity by the phosphorylation system, which may be particularly pronounced at maximum ETS capacity supported by convergent NS-electron supply. Uncoupling exerted only a slight effect on NS-pathway capacity. The median excess *E-P* capacity factor was 0.03 (0.02-0.06) and 0.03 (0.01- 0.07) in 32D cells, with or without IL-3 respectively. Inhibition of CI by rotenone in the presence of succinate induces the S-pathway control state (Fig. 4A). S-pathway ETS capacity was not reduced significantly following 8 h IL-3 withdrawal (Fig. 4C).

The OXPHOS coupling efficiency, 1-*L/P*, was 0.86 (0.79-0.90) and 0.82 (0.72-0.89), respectively, in control and IL-3 deprived 32D cells measured in the N-pathway control state with PGM (Fig. 4). This indicates a high degree of coupling in both cell types. By comparison, the ETS coupling efficiency, 1-*L/E*, was 0.86 (0.79-0.90) and 0.82 (0.72-0.89) in intact 32D+IL3 and 32D-IL3 cells, respectively, 0.84 (0.79-0.90) in 32Dv-RAF+IL3 cells, and 0.91 (0.81-0.99) in 32Dv-RAF-IL3 cells (Fig. 1C). Taken together, no uncoupling occurred prior to apoptosis, and comparable estimates of high coupling efficiency were obtained in permeabilized and intact cells, thus indicating that opening of the mt-permeability pore was not implicated. Similarly, in human leukemia cells, a high and constant mitochondrial coupling state was observed in the early phase of apoptosis ^34^.

### Complex IV is the main respiratory control variable of mitochondrial injury in 32D cells

Cytochrome *c* oxidase (Complex IV; CIV) is the terminal enzyme in the mitochondrial electron transfer system. Measurement of the enzyme activity of this single step in the pathway of oxygen consumption constitutes a key element in OXPHOS analysis ^43^. After inhibition of Complex III by antimycin A, residual oxygen consumption (ROX) was close to zero. Ascorbate and *N,N,N’,N’*-tetramethyl-*p*-phenylenediamine (TMPD) were then added to evaluate CIV activity corrected for autoxidation of ascorbate and TMPD (chemical background, chemBG; Fig. 4A). A significant decrease of CIV activity was observed in 32D-IL3 cells compared to control 32D+IL3 cells, similar to the decline of NS-pathway capacity (Fig. 4C). Therefore, it was of interest to determine the threshold level of CIV capacity as a potential basis to explain the decrease in cell respiration and NS-pathway flux. Cyanide titrations resulted in hyperbolic inhibition of CIV and NS-pathway flux (Fig. 5A). The threshold plot displays normalized NS-pathway flux as a function of CIV inhibition (Fig. 5B). Of note, the flux control ratio of NS-OXPHOS capacity (NS/NSref) is plotted as a function of CIV inhibition in the ETS state (1- CIV/CIVref; Fig. 5B). This is based on the result that *P* was nearly equivalent to *E* (*P/E*=0.97±0.01 SD in +IL3 cells, *P/E*=0.97±0.02 SD in -IL3 cells; Fig. 4C) in the NS(PGMS) pathway. Therefore, cyanide titrations were performed in the NS*P* state which provided higher experimental stability of flux compared to the NS*_E_* state. The excess CIV-NS capacity was essentially zero, as concluded from the fact that any inhibition of CIV caused a directly proportional inhibition of NS-pathway flux. The threshold for inhibition of CIV is defined as the intercept of the slope. An intercept of 1.0 illustrates the lack of apparent excess capacity with reference to convergent NS-electron flow (CIV0/NS_ref_; Fig. 5B). The consequences of zero excess CIV-NS capacity on interpreting the mt-flux control pattern (Fig. 5C) in 32D-IL3 cells are analyzed in the discussion.

**Fig. 5:**
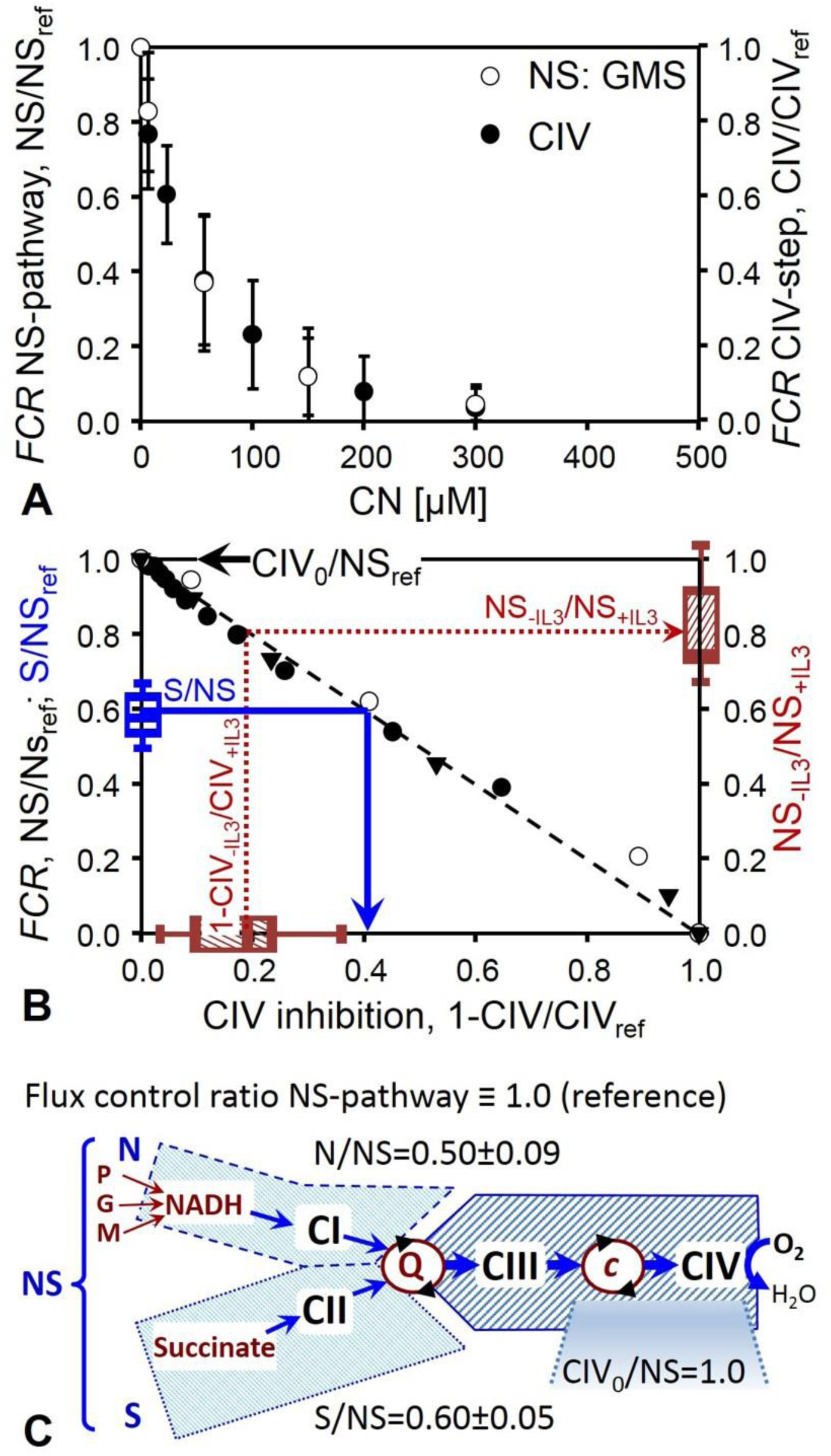
Cyanide titration and apparent excess CIV capacity in permeabilized 32D cells. (A) Permeabilized +IL3 control cells were titrated with cyanide in the presence of glutamate, malate&succinate and ADP (NS; open symbols). The flux control ratio, *FCR*, of the NS-pathway is expressed relative to the reference flux at zero inhibitor concentration, NS/NS_ref_. Using the same titration steps as for the pathway flux, cyanide titrations of CIV enzyme velocity were performed after inhibition of CI and CIII (rotenone and antimycin A) and stimulation of CIV by ascorbate&TMPD and uncoupler (CIV; filled symbols). The *FCR* of the single step is expressed relative to the reference CIV velocity at zero inhibitor concentration, CIV/CIV_ref_. (B) Threshold plot for apparent excess CIV capacity, relating the relative pathway flux through the electron transfer system as a function of relative inhibition of the single enzymatic step of CIV at identical cyanide concentrations. The CIV_0_/NS flux ratio is calculated as the intercept of the *Y*-axis at zero CIV inhibition (CIV_0_) of a linear regression, presenting a zero threshold with an intercept of 1.0 (dashed line). Therefore, the apparent excess capacity of CIV, *j*_ExCIV_ = CIV_0_/NS −1, equals zero (for comparison, see 43). Preserved S-pathway capacity is fully compatible with the decline of CIV activity after IL-3 withdrawal, as shown by the flux control ratio for the S-pathway in controls, S/NS (left *Y*-axis; from Fig. 4C), projected onto a threshold of relative CIV inhibition of 0.4 (full line, arrow to relative CIV inhibition), which is above the observed CIV-IL3 inhibition. CIV inhibition in the -IL3 group relative to +IL3 controls is plotted on the *X*-axis (1 - CIV_-IL3_/CIV_+IL3_; from Fig. 4C). The NS-pathway capacity in the -IL3 group relative to +IL controls is plotted on the right *Y*-axis (NS_-IL3_/NS_+IL3_; from Fig. 4C). The decline of CIV-IL3 capacity thus explains the proportional decline of NS-pathway capacity (dotted line, arrow to NS_-IL3_/NS_+IL3_). Results are means ± SD (*N*=3 cultures distinguished by different symbols, with duplicate experiments on each culture). (C) Schematic representation of NADH-linked and succinate-linked (N and S) pathways converging at the Q-junction (Coenzyme Q), and corresponding N/NS and S/NS flux control ratios in 32D+IL3 control cells.

## Discussion

In 32D cells, prolonged growth factor (IL-3) withdrawal results in growth arrest and subsequent apoptosis. After about 12 to 15 h of IL-3 deprivation, cells accumulate in the G1 phase, mitochondrial ROS production and mitochondrial Ca^2+^ levels are increased and the cells become irreversibly committed to death at a time point of 16 h ^27, 32^. Cell death and the preceding perturbations of mitochondrial Ca^2+^ and ROS homeostasis can be significantly delayed by oncogenic v-RAF ^27^. The objective of our investigation was to evaluate mitochondrial respiration before the onset of apoptosis. After 8 h IL-3 withdrawal, viability and cell cycle distribution of the 32D cells were not affected, but mitochondrial respiratory capacity was decreased.

To gain insight into the possible mechanisms underlying the reduction of respiratory capacity in IL-3 starved 32D cells, we tested the hypothesis that the change in mitochondrial respiration was caused by a general decrease in mitochondrial content or by a switch from aerobic to glycolytic pathways. In specific tumors, a mitochondrial injury causing an increased dependence on glycolysis for ATP production is well documented (Warburg effect, see reviews ^35-39^). We therefore assayed citrate synthase (CS), an enzyme of the TCA cycle, as a marker of mitochondrial content and lactate dehydrogenase as a glycolytic marker. The results show that the decrease in respiration cannot be explained by an enzymatic switch to glycolysis, since the activities of the two marker enzymes varied in direct proportion, without a significant change following IL-3 withdrawal. Similarly, succinate-supported respiratory capacity did not decline significantly in the 32D-IL3 cells compared to controls. In contrast, N- and NS-pathway capacity and CIV-activity were decreased in 32D cells after 8 h IL-3 withdrawal. A simple diminution of mitochondrial content would affect all mitochondrial pathways at the same proportion, without any change of the pattern of mitochondrial respiratory control. Thus 32D cells show a different pattern compared to leukemia cells where alteration of respiratory capacity and enzyme activities per cell are mainly caused by changes in cell size and mitochondrial content per cell, occurring upon cell cycle arrest triggered by apoptotic agents in the early phase of apoptosis ^34^.

Our data exclude cell size and mitochondrial content as explanations of the decrease of respiration in intact 32D cells after IL-3 withdrawal. Two other possible explanations for the decrease in ROUTINE respiratory activity are a shift to a metabolic state with lower energy demand, or mitochondrial injuries. Inhibition of intact cell respiration by the CI inhibitor rotenone (Fig. 3) is frequently thought to provide a diagnostic measure of electron flow through the NADH-pathway, when rotenone suppresses most of the respiratory activity ^24^. This interpretation ignores the fact that inhibition of CI simultaneously prevents formation of succinate in the TCA cycle, since succinate production is stopped when NADH cannot be oxidized by CI. Without cytosolic sources of succinate, therefore, rotenone inhibition of cell respiration does not exclusively inhibit electron flux through CI, but simultaneously suppresses the succinate-pathway flux by redox-coupling in the TCA cycle.

In the N-pathway, pyruvate, malate and glutamate dehydrogenases reduce NAD+ to NADH, feeding electrons into CI. Electrons are further transferred through CIII and CIV to oxygen, the final electron acceptor (N; Fig. 5C). CI, CIII and CIV are frequently arranged as supercomplexes ^40, 41^. When succinate is provided, electrons are transferred though succinate dehydrogenase (CII) to CIII and CIV (S; Fig. 5C). N- and S-pathway capacities were 0.50 and 0.60, respectively, of convergent NS-pathway capacity, thus exerting an almost completely additive effect on the combined pathway (NS; Fig. 5C). When measured separately, pathway control resides upstream of the Q-junction, and downstream catalytic capacities are in excess (Fig. 5C). When Complex IV is in excess, pathway flux does not decrease proportionally to inhibition of CIV activity due to a threshold effect. In most types of mitochondria, CIV is in excess of electron transfer pathway capacity, even at maximal upstream electron supply by the convergent NS-pathway ^42, 43^. In macrophage cells the apparent excess CIV capacity, (CIV-NS)/NS, is 0.46 ^44^. Similarly, in a variety of intact human cell lines, the apparent excess CIV capacity in intact uncoupled cells varies between 0.00 and 0.40 ^45-47^. In 32D cells, the excess CIV-NS capacity was practically zero. This suggests that CIV is the main respiratory control variable of mitochondrial injury in 32D cells. The observation of a preserved S-pathway capacity (Fig. 4C) does not contradict this conclusion, but provides evidence of a significant excess CIV capacity with respect to the S-pathway. The corresponding CIV threshold increases as the S/NS flux control ratio declines ^43^. Since the S/NS flux control ratio was as low as 0.6 (Fig. 5C), S-pathway capacity is not expected to decline as long as inhibition of CIV does not exceed 40% (Fig. 5B). An exclusive defect of CIV, however, cannot explain the significant decline of N-OXPHOS capacity, since the N/NS flux control ratio was even lower than S/NS, indicating an additional defect upstream of the Q-junction or in the CI+CIII+CIV supercomplex (Fig. 5C). In physiological states with convergent NS-electron flow, the apparent excess capacity of downstream electron transfer is lower and flux control is shifted towards CIII and CIV.

Cytochrome *c* normally resides in the inner mt-membrane and is an essential component of the respiratory system, responsible for electron transfer from CIII to cytochrome *c* oxidase ^48^. Cytochrome *c* is an important apoptogenic factor, since its release from the mitochondrial intermembrane space initiates caspase activation which leads to apoptotic cell death ^49-52^. The respirometric stimulation by addition of exogenous cytochrome *c* is an indirect method to show permeability of the outer mt-membrane and release of cytochrome *c* to the cytosol. In the 32D cells deprived of IL-3 for 8 h, cytochrome *c* release was detected to a small extent, without significant increase in cell death or uncoupling. Mitochondria remained highly coupled even after IL-3 withdrawal, as shown by high OXPHOS coupling efficiencies in the range of 0.72 to 0.90 measured in the N-pathway control state with PGM. This agrees with results on leukemia cells undergoing apoptosis ^34^. Expression of v-RAF residing in the outer mitochondrial membrane had also been shown previously to protect 32D cells ^53^, human leukemia cells ^34^ and fibroblasts ^8^ by preventing cytochrome *c* release and subsequent activation of caspases ^53^. A possible explanation for the RAF-induced inhibition of cytochrome *c* release in growth factor starved cells is the formation of a complex with the voltage-dependent anion channel (VDAC or porin; ^53^ a mitochondrial protein involved in the exchange of metabolites for oxidative phosphorylation ^54^). The new complex formed *in vivo* blocks reconstitution of VDAC channels in planar bilayer membranes *in vitro* ^53^. It is surprising, however, that cytochrome *c* release was detected at this early time point, when the 32D cells are still not presenting any increase in mortality and can fully recover if they are provided again with IL-3. These findings suggest that the cells deprived of IL-3 may have become more sensitive and experience some mitochondrial membrane fragility.

Our study provides novel insights into a key pathway involved in malignant transformation. Mitochondrial dysfunction is triggered by growth factor limitation, which is a critical step in cancer progression, when the increased size of the growing tumor affects the diffusion of oxygen and nutrients. The viral oncogene v-RAF has been discovered ^55^ long before mutations were detected in human BRAF ^56^ and CRAF ^1^. The murine homolog BRAF, therefore, might currently be considered as the most relevant oncogene. However, many key findings regarding signaling by oncogenic RAF have been initially obtained with v-RAF and were confirmed later with oncogenic variants of BRAF and CRAF. This suggests that the effects of activated RAF on mitochondrial physiology plays out in human tumors carrying mutant forms of BRAF or CRAF.

Our results demonstrate that IL-3 withdrawal severely compromises mitochondrial respiratory function in a fashion that is suppressible by v-RAF. This suggests a direct link between the key mitogenic and survival kinase CRAF and mitochondrial energy homeostasis. A pro-survival role of mitochondrial respiratory competence is recognized in various types of cancer cells ^57-60^, indicating that - in contrast to a cancer-specific Warburg effect – changes in mitochondrial respiratory control play a critical role in determining ‘positive’ or ‘negative’ fates in both normal and transformed cells. Taken together, these studies suggest that a paradigm shift is required in cancer biology, to avoid biases in interpretation of mitochondrial changes as Warburg-type dysfunctions and open our views to alternative concepts on compensatory or ‘adaptive’ responses of mitochondrial function in the context of cell metabolism, proliferation, and cell death. The decline of mitochondrial respiratory capacity comprised an early event in the pathway to apoptosis after growth factor withdrawal, before the onset of inactivation of the main signaling effectors of the IL-3 receptor. This time course points to a primary role of mitochondrial respiratory function in apoptosis of these cells.

## Materials and Methods

### Cell culture

Experiments were performed with the parental promyeloid interleukin-3 (IL-3)- dependent cell line 32D and with 32Dv-RAF cells expressing activated CRAF ^32, 61^. Cells were grown in RPMI 1640 (PAA Laboratories Pasching, Austria) supplemented with fetal calf serum (10%, heat-inactivated 45 min at 56 °C), 100 U of penicillin (100 U·ml^−1^; Invitrogen, Carlsbad, CA), streptomycin (100 U·ml^−1^; Invitrogen, Carlsbad, CA), L-glutamine (2 mM; Invitrogen, Carlsbad, CA) and WEHI cell-conditioned medium (15%). Every three days cells were split at a ratio of 1:10 and refed with fresh medium. Cell viability was determined by trypan blue exclusion assay and cells were used in experiments only when the viability was above 97%. For starvation experiments cells were washed three times in IL-3 free RPMI 1640 and resuspended in RPMI 1640 without IL-3 at a cell density of 0.5·10^6^ cells·ml^−1^ and incubated for 8 h (Fig. 1A). Measurement of respiration in intact cells was performed in RPMI (density 1·10^6^ cells·ml^−1^). For the cell permeabilization test and the measurement of respiration in permeabilized cells, cells were rinsed twice with respiration medium MIR05 (110 mM sucrose/ 60 mM K-lactobionate/ 0.5 mM EGTA/ 1 g·l^−1^ BSA fatty acid free/ 3 mM MgCl2/ 20 mM taurine/ 10 mM KH2PO4/ 20 mM K-HEPES/ pH 7.1 ^62^), and diluted to a concentration of 2·10^6^ cells·ml^−1^ in MiR05.

### Cell size determination

Cell volume was analysed in a CASY1 TT cell counter and analyser system (Schärfe System, Reutlingen, Germany) which combines the techniques of particle measurement and pulse area analysis signal processing. Cell suspensions were diluted in isotonic CASY Ton^®^ solution (100 μl in 10 ml) and introduced into the sensor through a capillary. Every cell is recorded with a frequency of 10^6^ evaluations per second during the pulse area analysis while passing through the sensor and the individual signals are analysed according to form, width, range and time frame. Events in the range of 6-25 μm diameter were included in the calculation.

### Cell cycle analysis

Cells were washed with 3 ml ice cold PBS and the pellet was resuspended in 500 μl ice cold PBS and 5 ml cold absolute ethanol added dropwise. Samples were centrifuged and supernatant was discarded. After a final wash with 5 ml ice cold PBS containing 1% BSA, samples were resuspended in 250 μl DNA staining buffer containing sodium citrate (1 mg·ml^−1^), Triton-X 100 (150 μl·ml^−1^), propidium iodide (0.1 mg·ml^−1^) and Ribonuclease A (0.02 mg·ml^−1^) and incubated at 37 °C for 30 min. Propidium iodide fluorescence was measured using linear FL2 channel in a FACSCalibur flow cytometer (Beckton Dickinson) and analyzed using Cellquest software. Resulting peaks corresponded with cell cycle states A, G1, S and G2 ^63^. Areas were set to calculate the four peaks in terms of percent of gated events and mean fluorescence intensity.

#### High-resolution respirometry

Cell survival was assayed before each experiment by staining with trypan blue. Cell respiration was measured at 37 °C with the Oxygraph-2k (Oroboros Instruments, Innsbruck, Austria) in 2-ml chambers at a stirrer speed of 750 rpm. Data acquisition and real-time analysis were performed using the software DatLab (Oroboros Instruments, Innsbruck, Austria). Automatic instrumental background corrections were applied for oxygen consumption by the polarographic oxygen sensor and oxygen diffusion into the chamber ^64^.

#### Intact cells

The protocol for the respiration in intact cells is illustrated in Fig. 1B. Respiration of intact cells was measured in culture medium RPMI (cell density c. 1 · 10^6^ cells·ml^−1^) under cellular routine conditions (ROUTINE). After inhibition of ATP-synthase with 2 μg·ml^−1^ oligomycin, respiration declined to the resting or leak-compensating state (LEAK, *L*). Uncoupling with stepwise titration to an optimal concentration of the protonophore carbonyl cyanide p-(trifluoromethoxy) phenylhydrazone, FCCP, induced maximal respiration as a measure of electron transfer system capacity (ETS, *E*). Residual oxygen consumption (ROX) was obtained after inhibition of CI and CIII with 0.5 μM rotenone and 2.5 μM antimycin A. Fluxes in all states were corrected for ROX and expressed per million cells.

FCCP titration was performed using the Titration-Injection-microPump TIP2k (Oroboros Instruments, Innsbruck, Austria). Two Hamilton syringes fitted with 27 mm needle length and 0.09 mm needle inner diameter were mounted on the TIP2k for simultaneous titration into the two chambers of the Oxygraph-2k. A 10 mM stock solution of FCCP was filled into the Hamilton syringes and 0.1 μl (0.5 μM) step titrations were performed at a titration speed of 20 μl·s^−1^. The optimum FCCP concentration for maximal respiratory oxygen flow varied from 3 to 4 μM. Since the uncoupler and inhibitors were dissolved in ethanol, controls were treated with corresponding titrations of ethanol, and no effect was detected.

#### Plasma membrane permeabilization

The optimum digitonin concentration for selective permeabilization of the plasma membrane was determined in 32D and 32Dv-RAF cells grown in the presence of IL-3 or after 8 h IL-3 withdrawal. The cell density for the permeabilization test was 2 million cells per ml. ROUTINE respiration as supported by endogenous substrates was measured in mitochondrial respiration medium MIR05. After addition of rotenone (0.5 μM), succinate (10 mM), and ADP (2.5 mM), digitonin was titrated in steps of 1, 2, 3, 4, 6, 10 and 20 μg·ml^−1^ ^65^ (Fig. 3).

#### Permeabilized cells

After adding inact cells (ce) for measurement of ROUTINE respiration in MIR05 (1ce) and permeabilization of cell membranes with optimal digitonin concentration (10 μg·ml^−1^), the following substrates, uncoupler and inhibitors were titrated (final concentration in the chamber): glutamate (G; 10 mM), malate (M; 5 mM), and pyruvate (P; 5 mM) as NADH-linked substrates (1PGM); ADP (1 mM), cytochrome *c* (*c*; 10 μM; 2D*c*); succinate (S; 10 mM) as CII-substrate (3S), FCCP (4U; optimum concentration 3 to 4 μM); rotenone (5Rot; 0.5 μM) and antimycin A (6Ama; 2.5 μM) as CI and CIII inhibitors; ascorbate (0.5 mM) and TMPD (2 mM) as CIV-linked substrates. Chemical background corrections were applied to account for autoxidation of TMPD and ascorbate (in the presence of 280 U·ml^−1^ catalase to prevent accumulation of hydrogen peroxide ^66, 67^).

#### Enzyme activity and protein assays

For determination of enzyme activities and protein content, 300 μl cell suspension was pipetted from the Oxygraph-2k chamber at the end of the experiments, snap-frozen in liquid nitrogen and stored at −80 °C until further analysis. Enzyme activities in each samples were assayed in duplicates at 30 °C and expressed in units of U·10^−6^ cells. U is one μmol of substrate transformed per minute. Citrate synthase activity was measured at 412 nm recording the linear reduction of 0.1 mM 5,5’dithiobis-2-nitrobenzoic acid (*ε*_412_: 13.6 ml·cm^−1^ · μmol^−1^) in the presence of 0.31 mM acetyl-CoA, 0.5 mM oxalacetic acid, 0.1 M Tris/HCl, 50 μM EDTA, 5 mM triethanolamine hydrochloride (pH 8.1; ^68, 69^). Lactate dehydrogenase (LDH) activity was measured at 340 nm recording the linear decrease of absorbance of 0.3 mM NADH 340 nm (*ε*_340_ = 6.22 ml·cm^−1^ · μmol^−1^) in the presence of 0.25% Triton X-100 (Serva, Vienna, Austria) and 10 mM pyruvate (Fluka, St. Louis, MO, USA, 0.1 M Tris/HCl buffer, pH 7.1 ^68^). Protein concentration was determined with the Lowry method (Bio-Rad protein assay kit, Richmond, CA).

#### Reagents

All chemicals were from Sigma-Aldrich unless specified.

#### Statistical analysis

The data are presented as medians (min-max). Comparing means with the medians showed differences <5% without affecting any statistical conclusions. Standard t-tests were used to calculate significance levels between the treatments (with or without IL-3). To determine the effects of addition of cytochrome *c*, or uncoupler, t-tests for dependent samples were used. Significance was considered at *P* <0.05.

## Author Contributions

HL and PS performed the respirometric experiments. CD and MW performed the cell size and cell cycle determinations. EG and JT conceived the project and provided supervision. HL and EG analyzed the data, and wrote the manuscript. JT was responsible for the cell culture and co-wrote the manuscript. EG revised the manuscript.

## Acknowledgments

We thank Tina Goebel for providing technical support for cell culture and Michaela Schneider for performing enzyme activity measurements. This work was supported by a grant from the Austrian Science Foundation, project MCBO ZFW011010-08, the Austrian Cancer Society/Tyrol, and K-Regio project MitoFit (EG and Oroboros Instruments, Innsbruck, Austria). HL was supported by postdoctoral fellowships from the ‘Fonds de Recherche sur la Nature et les Technologies’ (Quebec government, Canada) and the National Sciences and Engineering Council of Canada. Contribution to COST Action CA15203 MITOEAGLE.

